# Spatial transcriptomic analysis of adult hippocampal neurogenesis in the human brain

**DOI:** 10.1101/2023.05.02.539011

**Authors:** Sophie Simard, Reza Rahimian, Maria Antonietta Davoli, Stéphanie Théberge, Natalie Matosin, Gustavo Turecki, Corina Nagy, Naguib Mechawar

## Abstract

Using spatial transcriptomics and FISH, we detected very few cells expressing neural stem cell- and proliferation-specific genes in the human dentate gyrus (DG) from childhood to middle age. However, we observed at all ages a significant number of DG cells expressing the immature neuronal marker *DCX*. Across ages, the majority of these cells displayed an inhibitory phenotype, while the remainder were non-committed or excitatory in nature.

## MAIN

Occurrence of adult hippocampal neurogenesis (AHN) in the subgranular zone (SGZ) of the human dentate gyrus (DG) was suggested for the first time 25 years ago^1^. AHN has been extensively investigated, but mainly with immunohistochemical approaches, yielding at times highly discordant conclusions^2-9^. However, recently, there has been a proliferation of studies using single-cell and single-nucleus RNA sequencing to examine AHN at the transcriptomic level^10-14^. While these methods provide transcriptomic signatures for different cell types detected in the DG, they lack spatial resolution.

To better contextualize the spatial profiles of neurogenesis markers in the human hippocampus, we used the 10x Genomics Spatial Gene Expression platform on frozen hippocampal sections from young and middle-aged neurotypical adults (Supplementary Table 1). We performed unsupervised clustering on the spots in each DG section using the R package BayesSpace^15^ (Fig. 1a; Extended Data Fig. 1a-c). The morphology of the DG in each sample was validated using both H&E staining and fluorescent *in situ* hybridization (FISH) with a probe directed against *PROX1*, a DG enriched gene^16^ (Fig. 1b). We next assessed the expression levels of canonical cell type markers for each cluster and observed their concordance with the expected spatial location of the clusters within the hippocampus (Fig. 1c). Interestingly, a subset of spots in Cluster 5 (56.3% of spots) located within the granule cell layer (GCL) of each section, expressed the inhibitory neuronal marker *GAD1*, alongside high levels of expression for the excitatory neuronal marker *SLC17A7* (Supplementary Table 2).

**Figure 1.**
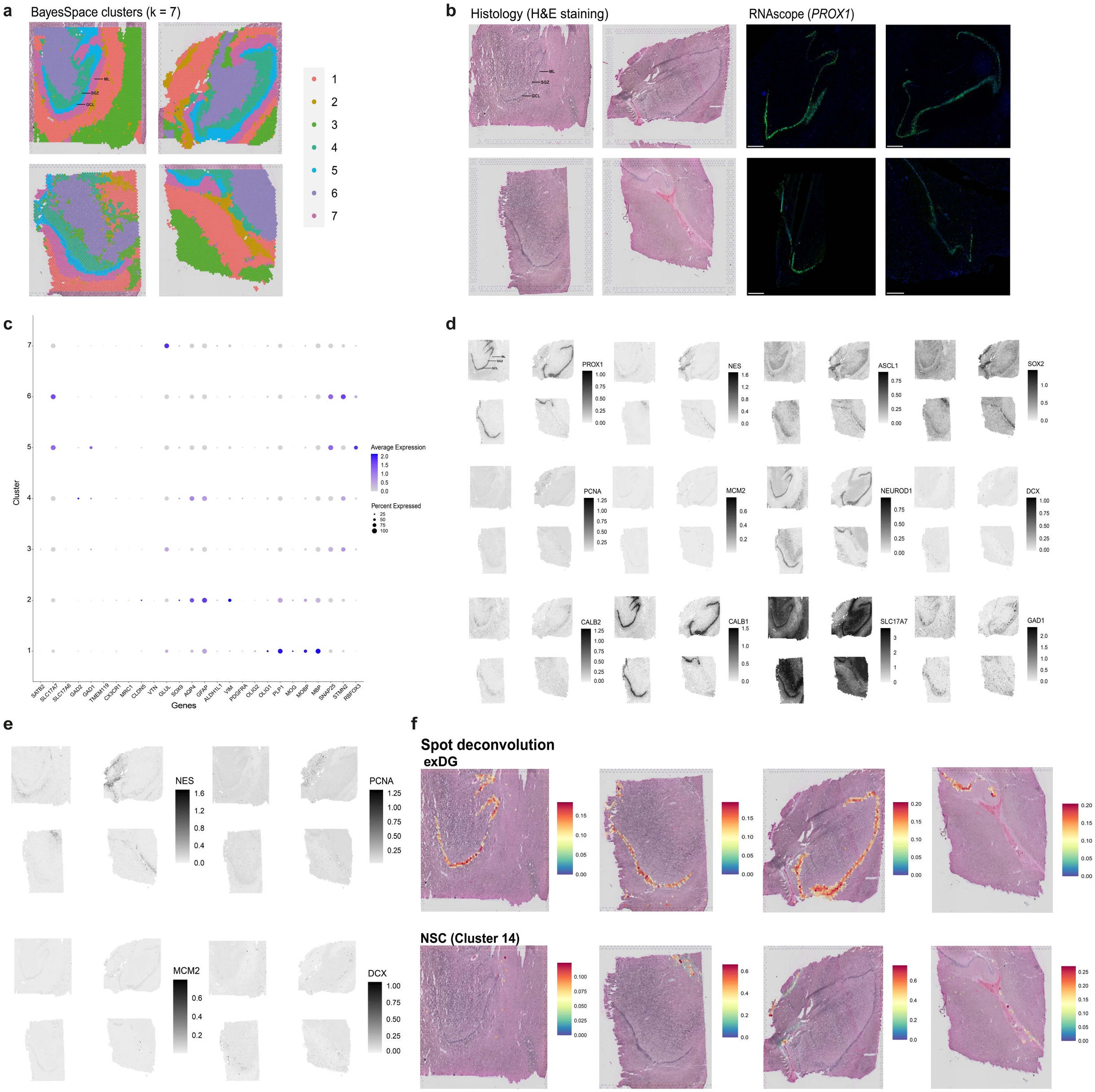
Detection of neurogenesis markers in Visium Spatial Gene Expression data of the adult human hippocampus and spatial reconstruction of a human hippocampal single-nucleus RNA sequencing dataset. **(a)** Plots showing the seven clusters (k=7) generated with the R package BayesSpace’s t-distributed error model algorithm (v. 1.4.1) on each section (n=4). The layers of the DG are labeled on the top left plot and the lines are labeling the SGZ, GCL and ML of the DG. Sections on the left are from middle-aged males (n=2, mean age=42.5 years), and sections on the right are from young adult males (n=2, mean age=23.5 years old), **(b)** Validation of DG morphology in each section with an H&E staining and RNAscope using a probe directed against the dentate-lineage marker *PROX1* (in green) with DAPI nuclear staining in blue. The layers of the DG are labeled on the top left plot and the lines are labeling the SGZ, GCL and ML of the DG. Scale bars for RNAscope scans = 800 μm**. (c)** Dotplot visualization of the scaled average expression of canonical cell type markers in each BayesSpace cluster in all sections. *SATB2, SLC17A7, SLC17A6* (excitatory neurons); *GAD1, GAD2* (inhibitory neurons); *TMEM119, CX3CR1, MRC1* (microglia); *CLDN5, VTN* (endothelial cells); *GLUL, SOX9, AQP4, GFAP, ALDH1L1, VIM* (astrocytes); *PDGFRA, OLIG1, OLIG2* (oligodendrocyte precursor cells); *PLP1, MOG, MOBP, MBP* (oligodendrocytes); *SNAP25, STMN2, RBFOX3* (neurons). The size of the dot corresponds to the percentage of spots within a cluster enriched for the marker. The colour of the dot represents the scaled average expression levels of the marker across spots within a cluster. **(d)** Log-normalized expression of neurogenesis markers spatially plotted onto each section at the sub-spot level using BayesSpace. Higher values in the scale correspond to higher expression levels, whereas lower values reflect lower expression levels. *NES, SOX2, ASCL1* (neural stem cells); *PCNA, MCM2* (proliferative cells); *NEUROD1* (neuroblasts); *DCX, CALB2* (immature granule neurons); *CALB1* (mature granule neurons); *PROX1* (dentate lineage cells); *SLC17A7* (excitatory neurons); *GAD1* (inhibitory neurons). **(e)** Enlarged sub-spot level plots showing *NES*, *PCNA*, *MCM2* and *DCX* log-normalized expression on each section. **(f) Upper panel:** plots representing the spatial reconstruction of the exDG (dentate granule cell) cluster from Habib *et al.* (2017) with the R package SPOTlight (v. 0.1.7). **Lower panel:** plots representing the spatial reconstruction of the NSC (neural stem cell) cluster from Habib *et al.* (2017) with SPOTlight. The coloured scale corresponds to the proportions of cell type markers from the single-nucleus clusters represented in each spot. SGZ, subgranular zone; GCL, granule cell layer; ML, molecular layer.

We next sought to spatially map markers specific to neural stem cells, proliferative progenitor cells, neuroblasts, immature granule neurons, mature granule neurons, and cells from the dentate lineage on each section at the sub-spot level (Fig. 1d). We observed *NEUROD1* expression in neuronal clusters, mainly the *PROX1*-enriched GCL, which corroborates previous findings of Prox1^+^NeuroD1^+^ immunolabeled cells in the rodent DG^17^. While many of the neurogenesis markers were expressed in either the SGZ or GCL, the proliferative markers *PCNA* and *MCM2* displayed very low expression in all samples (Fig. 1e). Moreover, *DCX* showed dispersed expression within the DG, and was also detected in hippocampal regions outside of the DG (Fig. 1e). Of note, we expected to find neural stem cell markers outside the DG since there are similarities between the transcriptomic profiles of human neural stem cells and astrocytes^18^. *NES* was also expressed in sub-spots enriched for oligodendrocyte precursor cell (OPC)-specific markers (Fig. 1c,e), indicating possible similarities between the transcriptomic signatures of both cell types. Since we unexpectedly detected *GAD1* in our GCL cluster (Cluster 5), we assessed *GAD1*’s spatially-resolved expression within each section (Fig. 1d) and observed that the expression of the marker mapped to the DG.

Previously, Habib *et al*. (2017) identified a hippocampal cluster with a transcriptomic profile suggestive of neural stem cells^12^. To better identify this cluster, we spatially reconstructed it as well as their DG granule neuron (exDG) cluster with spot deconvolution^19^ (Fig. 1f). We observed that the spots located in the GCL had higher proportions (10%-20%) of the transcriptomic profile for the exDG cluster, whereas the predicted spatial location of the NSC cluster did not correspond to the SGZ, but rather to regions outside of the DG. These results further support the recent re-analysis of the dataset in which the authors re-examined the transcriptomic identity of the NSC cluster^20^ by performing gene set enrichment using a set of previously identified human ependymal cell markers^21^.

Altogether, these data reveal that neurogenesis markers are spatially resolved to cells in the DG. However, they also map to regions outside of the typical hippocampal neurogenic niche, which raises questions as to the specificity of these markers, especially immature neuronal markers, to AHN. The findings further confirm the importance of using multiple markers to characterize different neurogenic cell types in the human DG.

The controversy in the literature regarding the extent to which AHN occurs in the human brain can be attributed to a wide range of factors, including differences in the brain specimens analyzed^22-24^ and immunohistochemistry protocols used to identify neurogenic cell types^4,20^, specifically antibodies used to define neurogenesis in the human DG^23^. These differences have resulted in diverging conclusions regarding the presence of different neurogenic cells immunolabeled in the adult human hippocampus.

To investigate these cells at the transcriptomic level, we used multiplexed fluorescent *in situ* hybridization in the SGZ and GCL of one infant, one adolescent and six adult hippocampal samples (Supplementary Table 3). Using a combination of probes to identify neural stem cells, we found *NES*^+^*SOX2*^+^ cells also expressing (or not) *ALDH1L1* in very low numbers in the SGZ, and almost none in the GCL, of the adolescent and adult subjects (Fig. 2a), which is congruent with the findings of Boldrini and colleagues^2^. These presumed neural stem cells represented less than 1% of all SGZ cells in the adolescent and adults. We used *PCNA* and *MCM2* probes to estimate cell proliferation in adult DG but were unable to quantify their expression levels since one cell/DG section at most displayed a signal above threshold for either of these markers (Fig. 2b). The low detection levels of these markers are consistent with their predicted spatial feature mapping in Visium data (Fig. 1d).

**Figure 2.**
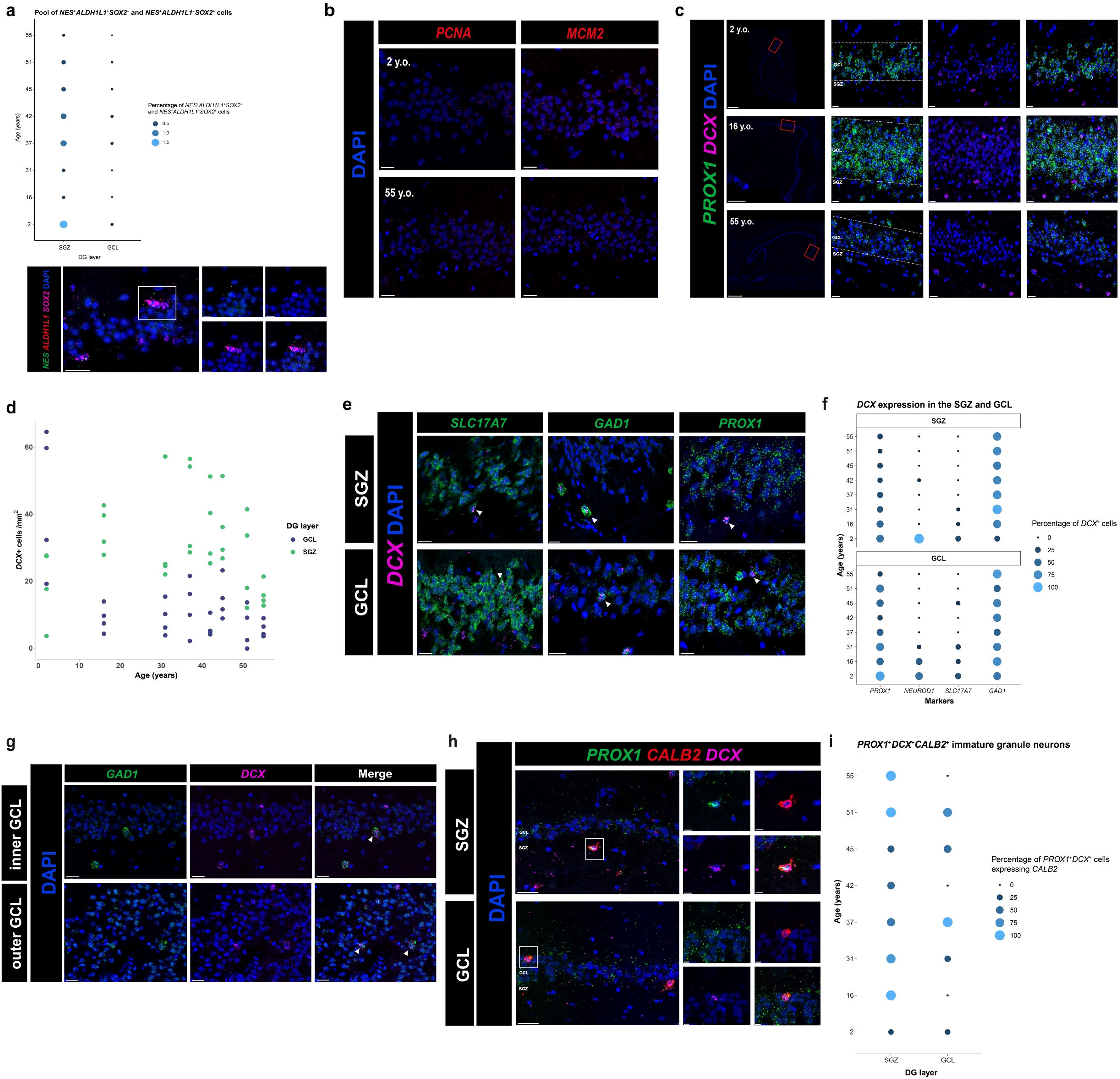
Distribution of neural stem cell-, proliferation- and immature neuron-specific genes and *DCX* expression combined with different cell type markers in the SGZ and GCL of the human DG. **(a)** Percentage of *NES*^+^*SOX2*^+^ expressing or not *ALDH1L1* cells in the SGZ and GCL of the whole DG across ages. Lower panel shows a *NES*^+^*SOX2*^+^*ALDH1L1*^+^ cell in the GCL of the infant DG (age=2 years old). Scale bars = 20 μm (scale bars for expanded insets = 10 μm). **(b)** *PCNA* and *MCM2* expression in the infant (age=2 years old) and adult dentate gyrus (age = 55 years old). Scale bars = 20 μm. **(c)** *DCX* expression detected in *PROX1*^+^ and *PROX1*^-^ cells in the DG of different age groups. Whole DG slide scanner images with scale bars = 1 mm (scale bars for expanded insets = 20 μm). Dotted lines delineate the SGZ and GCL of the DG. **(d)** Number of *DCX*^+^ cells per mm^2^ in the SGZ and GCL of the whole DG in infant (n=1), adolescent (n=1, age=16 years old) and adult subjects (n=6, mean age=43.5 years old). The graph shows values of four staining replicates per subject. **(e)** *DCX* expression detected in cells expressing different markers located in the SGZ and GCL. *PROX1* (dentate lineage); *NEUROD1* (neuroblasts); *SLC17A7* (excitatory neurons); *GAD1* (inhibitory neurons). Scale bars = 20 μm. **(f)** Percentage of *DCX*^+^ cells expressing *PROX1, NEUROD1*, *SLC17A7* and *GAD1* in the SGZ and GCL of the whole DG across ages. **(g)** *DCX*^+^*GAD1*^+^ cells located in the inner GCL (closer to the SGZ) and outer GCL (closer to the molecular layer). Scale bars = 20 μm. **(h)** *DCX*^+^ cells expressing *PROX1* and *CALB2* in the SGZ and GCL. Scale bars = 20 μm (scale bars for expanded insets = 5 μm). **(i)** Percentage of *PROX1*^+^DCX^+^ cells expressing *CALB2* and *DCX* in the SGZ and GCL of the whole DG across ages. White arrows point to double-positive cells. SGZ, subgranular zone; GCL, granule cell layer.

The low combined expression of *NES*, *SOX2* and *ALDH1L1*, as well as the low expression of *PCNA* and *MCM2* that we observed in all subjects suggests that proliferation in the DG is either mostly or entirely completed prenatally. These findings are consistent with previous reports suggesting that proliferation in the human DG drops significantly during infancy^7^, and human DG development occurs prenatally^25^.

To address the debate of DCX-immunolabeled cells in the DG^2, 3, 5-7, 9, 20^, we labeled *DCX* transcripts in the SGZ and GCL (Fig. 2c). *DCX*^+^ cells in the GCL decreased from an average of 44.1 (SD **±** 21.8) cells/mm^2^ in the 2 year-old subject to 8.9 (SD **±** 4.1) cells/mm^2^ in the 16 year-old subject, while *DCX*^+^ cells in the SGZ remained similar across all age groups (Fig. 2d; Extended Data Fig. 2a).

We next investigated the possible phenotypes of *DCX*^+^ cells detected in the AHN-associated DG layers by combining *DCX* with different cell type markers: *SLC17A7* (excitatory neurons), *GAD1* (inhibitory neurons), *NEUROD1* (neuroblast-specific)^17, 26^ and *PROX1* (dentate lineage)^16^ (Fig. 2e). Quantification of *SLC17A7* and *DCX* co-expression revealed a decrease in the percentage of *DCX*^+^ cells expressing *SLC17A7* in both DG layers in adults (Fig. 2f; Extended Data Fig. 2c,d). Strikingly, about half of *DCX*^+^ cells were found to express *GAD1* across all ages in the SGZ and the GCL, with cells located either in the inner GCL or the outer GCL, in the vicinity of the molecular layer (Fig. 2e-g; Extended Data Fig. 2c,d). Conversely, the decline in the percentage of *DCX*^+^*NEUROD1*^+^ cells, becoming barely detectable in adults, correlated with aging in both layers (Fig. 2f; Extended Data Fig. 2c,d). Finally, we found a decrease in the percentage of *DCX*^+^ cells expressing the dentate lineage marker in the adult subjects in both DG layers although this correlation only reached significance in the SGZ (Fig. 2e,f; Extended Data Fig. 2c,d). Our findings suggest that the majority of *DCX*^+^ cells in the adult human DG may display a GABAergic phenotype (average of 61% in the GCL and 75% in the SGZ), whereas only a minority of *DCX*^+^ cells were committed to a glutamatergic fate (average of 5% in the GCL and 1% in the SGZ). We also note that a significant subset of *DCX*^+^ cells did not express *SLC17A7* or *GAD1*, and therefore may represent immature neurons not yet committed to a particular phenotype. It is also possible that at least some of these *DCX*-expressing cells correspond to glial cells^7,20^, or to cells with multipotentiality^27^, such as OPCs^28^. Indeed, previous studies have reported *DCX* expression in cell types that are not part of the dentate lineage, including microglia^7^ and astrocytes^29^, in the human brain. Consistent with these findings, our results reveal that *DCX* transcripts are detected in *TMEM119*^+^ microglia (Extended Data Fig. 3a), and *ALDH1L1*^+^ astrocytes (Extended Data Fig. 3b) in the adult human DG.

To investigate the presence of a pool of immature granule neurons in the DG throughout normal aging, we used a combination of *PROX1*, *DCX* and *CALB2* (calretinin) probes (Fig. 2h) based on previous studies^2, 5, 30, 31^. Quantification of these probes in the SGZ and GCL revealed that *PROX1*^+^*DCX*^+^ cells expressing *CALB2* in the SGZ in adults were generally in majority (average of 73%) (Fig. 2i). We also detected transcripts of *STMN1,* a tubulin-depolymerizing protein, which is enriched in human immature granule neurons^14^, in *PROX1*^+^*DCX*^+^ cells located in both layers of the DG in infant and adult hippocampi (Extended Data Fig. 4a,b) and validated *STMN1* expression in the DG with our spatial transcriptomic data (Extended Data Fig. 4c).

Whether these immature granule neurons are newly generated or not remains unknown. However, we speculate that these immature neurons may have different origins, as previously stated in Seki (2020)^23^ and as follows: cells expressing immature neuronal markers may reflect (1) neurons generated prenatally that persist during adulthood by remaining in a dormant-like state, (2) neurons generated postnatally that are a result of minimal proliferation in the DG and remain in a prolonged state of immaturity, which corroborates the idea of a delayed maturation of immature dentate granule neurons, thought to occur in long-living and large-brained species^32-34^, and (3) neurons that are re-expressing immature neuronal markers according to the recent hypothesis of neuronal dematuration in the DG^23, 35^.

Previous reports have shown expression of *DCX* in non-neurogenic regions, such as the human amygdala^20,36-38^ and cerebral cortex^36^. These findings raise concerns regarding the validity of DCX as an appropriate proxy for adult brain neurogenesis. To further confirm DCX expression in non-neurogenic regions^20,36-38^, we first spatially mapped the expression of *DCX* and *PROX1* onto publicly available spatial genomic data of dorsolateral prefrontal cortex (DLPFC) sections^39^. *DCX* expression was visible in a few sub-spots in all four DLPFC sections, whereas *PROX1* is mainly expressed in sub-spots located in the cluster annotated as white matter (Fig. 3a). These results confirm that expression of these markers may be found outside of the neurogenic niche of the human DG, which is in agreement with our spatial transcriptomic findings (Fig. 1d). We also confirmed the presence of *DCX* expression in the DLPFC using FISH on frozen unfixed sections (Fig. 3b).

**Figure 3.**
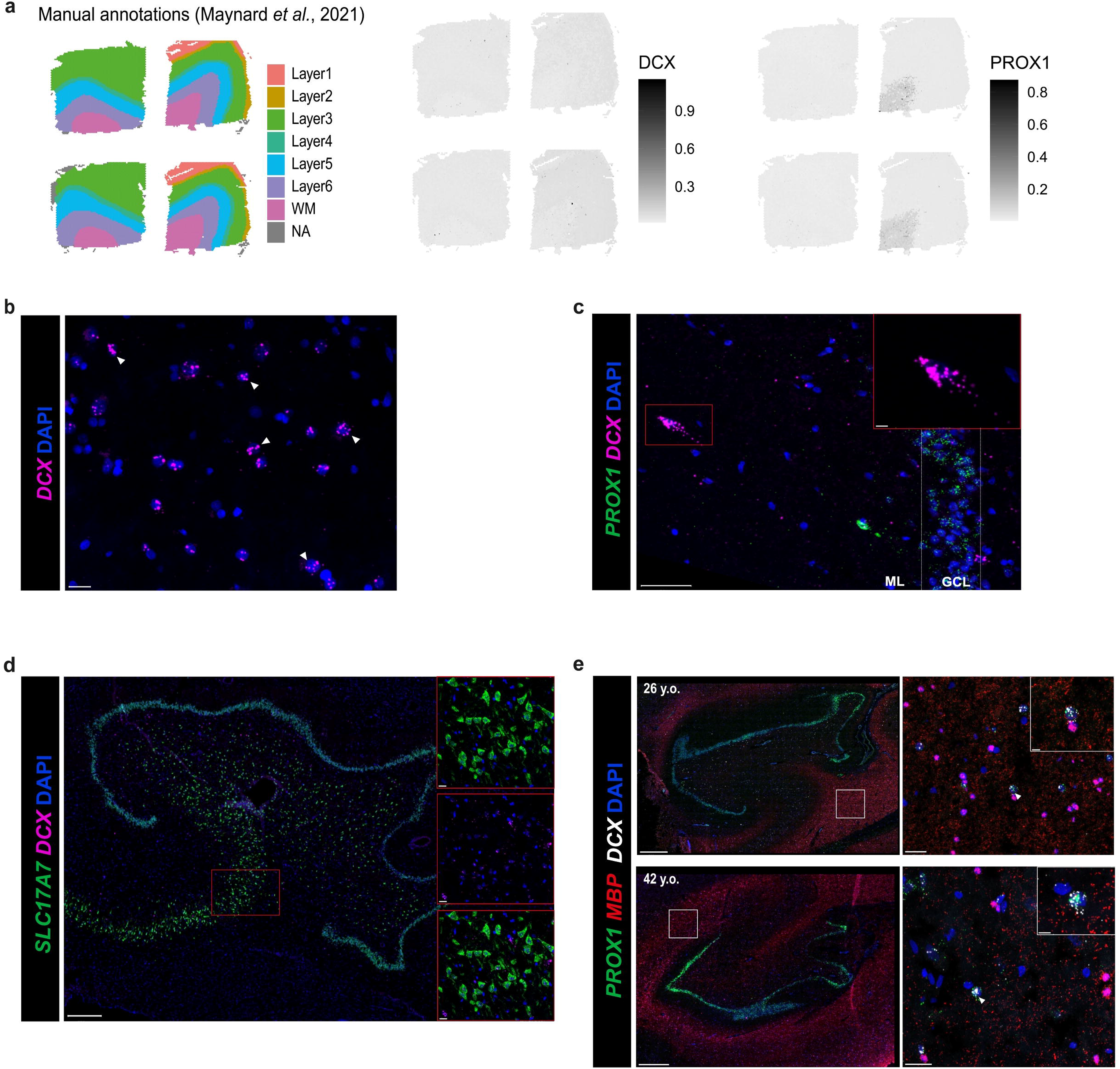
Specificity and distribution of *DCX* in the human brain. **(a) Left panel:** Manual annotations of samples 151671, 151672, 151673 and 151674 from Maynard *et al*. (2021) (n=4, 2M/2F, mean age=38.41 years old) plotted with the R package BayesSpace and showing the six cortical layers (Layers 1-6) and a white matter region (WM). **Middle and right panels:** Log-normalized expression of *DCX* and *PROX1* spatially plotted onto each section at the sub-spot level using BayesSpace. Higher values in the scale correspond to higher expression levels, whereas lower values reflect lower expression levels. **(b)** *DCX* expression with DAPI staining in a DLPFC section of a 26-year-old subject using RNAscope. Scale bars = 50 μm. White arrows point to *DCX*^+^ cells. **(c)** *DCX* expression detected in the ML of the DG of a 37-year-old subject. Scale bars = 50 μm and 10 μm (expanded inset). **(d)** *DCX* expression in cells located in the CA3 region of the hippocampus in a 51-year-old subject. Whole DG slide scanner image with scale bar = 500 μm. Higher magnification confocal images with scale bars = 20 μm. **(e)** *DCX* expression in *PROX1*^+^ cells located in regions highly expressing *MBP* in the adult human hippocampus. Whole DG slide scanner images with scale bars = 800 μm. Higher magnification confocal images with scale bars = 20 μm. SGZ, subgranular zone; GCL, granule cell layer; ML, molecular layer.

Additionally, we found *DCX*^+^ cells located in the molecular layer of the DG (Fig. 3c), a region where DCX-immunolabeled cells have been previously observed^11^. Finally, *DCX* expression was also detected in other hippocampal areas, such as CA3, although infrequently (Fig. 3d), and in *PROX1*^+^ cells from regions highly expressing the oligodendrocyte specific marker *MBP* (Fig. 3e). These observations suggest that multipotential cells, such as OPCs, expressing *DCX* could migrate out of the DG circuitry to reach their final destination and mature into the required cell type^28^, but this hypothesis requires further examination. Altogether, our data suggest markers commonly used to examine AHN may be insufficient to specifically define neurogenic cell types as they are expressed in brain regions not known to be neurogenic, at least in rodents.

The present study examines AHN in subjects from a wide age range and provides important new insights into the spatial expression of several neurogenesis markers as well as the possible phenotypes of *DCX*^+^ cells in the DG of the healthy human brain. Our findings indicate that the adult human DG exhibits, at most, very low levels of neurogenesis, due to the scarcity of neural stem cells and lack of expression of proliferative markers. The association between high brain complexity and low hippocampal neurogenesis in humans is a paradoxical observation that has been highlighted in previous studies^40, 41^.

While our data suggest that DG neurogenesis is not a significant phenomenon in humans postnatally, the small population of immature granule neurons in this region might play a more important role in driving hippocampal plasticity. Such cells have previously been suggested to contribute to the maintenance of a neurogenic reserve^42^, and to provide lifelong flexibility to the brain region, and act as a buffer against age-related hippocampal functional deficits^42, 43^. It is therefore possible that in the human DG, these immature granule neurons contribute substantially to preventing age-related brain damage^33^. Some experimental evidence supports this hypothesis in the adult rodent hippocampus^44, 45^, however, future work should investigate the relevance of this hypothesis in the adult human DG.

The expression of *GAD1* in *DCX*^+^ cells in the GCL was an unexpected finding. Although an earlier paper reported weak immunoreactivity for GAD1 in GCL neurons in the healthy human DG^46^, rodent studies have identified and characterized Gad1 expression in the GCL, and more precisely in the inner part of the layer adjacent to the SGZ^47, 48^, suggesting that Gad1^+^ cells displaying an immature phenotype in this layer are undergoing late neuronal differentiation^47^. Similarly, another study detected co-localization of DCX and GAD1 in DG granule cells from younger and older human subjects^49^. Thus, it is plausible that most of the *DCX*-expressing cells of the GCL that we identified are in a late granule cell differentiation stage since the number of *DCX*^+^ cells expressing *NEUROD1*, an early neuronal differentiation marker^27^, decreases in the DG of older individuals, while the number of *DCX^+^* cells expressing *GAD1* remains high. The present identification of GCL *GAD1*^+^*DCX*^+^ cells across all subjects confirms the presence of this subpopulation and warrants further investigation of their functional properties.

The scope of this study is limited to mRNA detection of various neurogenesis markers in the DG. Our spatial transcriptomic experiments using Visium Spatial Gene Expression, as well as our *in situ* hybridization studies were also performed on small sample sizes and only included male subjects. Despite these limitations, the present results also provide support for the hypothesis that the neurogenic capacity of the adult human DG may rely primarily on a reserve of immature granule neurons. Understanding the functional significance of the presence of a population of immature granule neurons during physiological aging can push forward future research on how DG neuroplasticity is altered in brain disorders.

## Methods

### Human post-mortem tissue

Frozen dentate gyrus (DG) tissue from well-characterized adolescent (n=1, age=16 years old), young adult (n=2, mean age=23.5 years old) and middle-aged males (n=2, mean age=42.5 years old), and frozen-fixed DG tissue from middle-aged males (n=6, mean age=43.5 years old) were obtained from the Douglas-Bell Canada Brain Bank (Montreal, Canada). In collaboration with the Quebec Coroner’s Office and with informed consent from next of kin, phenotypic information was obtained with standardized psychological autopsies. Presence of any or suspected neurological/neurodegenerative disorder signaled in clinical files constituted an exclusion criterion. Controls are defined with the support of medical charts and Coroner records. Toxicological assessments and medication prescription were also obtained. The samples used for spatial transcriptomic experiments were dissected from the anterior hippocampus and the body (close to the anterior pole) of the hippocampus, while all frozen-fixed samples used for RNAscope were dissected from the anterior hippocampus. The infant frozen dentate gyrus sample (age=2 years old) was provided by Dr. Nada Jabado’s laboratory from the The Research Institute of The McGill University Health Center. Detailed information on the samples used for spatial transcriptomic and fluorescence *in situ* hybridization experiments are presented in Supplementary Tables 1 and 3.

### Tissue processing and Visium data generation

Frozen tissue embedded in OCT were cryosectioned at -20 °C. Sections of 10 μm thickness were placed in the 6mm by 6mm fiducial frame capture area of the Visium Spatial Gene Expression Slides (catalog no. 2000233, 10x Genomics) to maximize tissue coverage. The tissue sections were next fixed in methanol at -20 °C for 30 minutes and stained with hematoxylin and eosin (H&E) according to the Visium Spatial Gene Expression User Guide (catalog no. 2000233, 10x Genomics). The tissue was permeabilized for 12 minutes based on tissue optimization time-course experiments. Brightfield images were taken using the Olympus VS120 slide scanner with a 10X objective and exported as low-resolution and high-resolution .tiff files. Libraries were prepared according to the Visium Spatial Gene Expression User Guide (CG000239) and loaded at 300pM. They were next sequenced on a NovaSeq 6000 System (Illumina) using a NovaSeq S4 Reagent Kit (200 cycles, catalog no. 20027466, Illumina), at very high sequencing depths ranging between 290-420 10×10^6^ read-pairs per sample. Sequencing was performed using the following read protocol: read 1: 28 cycles; i7 index read: 10 cycles; i5 index read: 10 cycles; and read 2: 90 cycles.

### Visium data processing

To process the sequenced data, the 10x Genomics analysis pipeline Space Ranger (spaceranger-1.1.0 version) was used to create FASTQ files which were mapped to the 10x Genomics GRCh38 reference human genome (GRCh38-2020-A) for gene quantification. QC metrics returned by this software are presented in Supplementary Table 1. The Space Ranger count pipeline was used to generate output files, including feature barcode matrices, Loupe Browser files for data visualization, data summary with images, metrics, and plots, downsampled input images, and spot barcode locations, for downstream analysis.

### Manual annotation

Each section was manually annotated on LoupeBrowser, the 10x Genomics Visium data visualization software, by assigning each spot to a specific region of the DG. The manual annotations were done based on the human anatomical atlas from the Allen Brain Atlas and validated by a blinded histology expert. Following manual annotation, seven clusters were defined on each section.

### Clustering accuracy comparison and unsupervised spot clustering

The raw Visium files for each sample (see Visium data processing) were read into R (v. 4.1.0)^50^ in a customized structure using the Seurat R package^51^ (V4) to keep them paired with the low-resolution histology images for visualization purposes. A quality check was first done with Seurat on each sample. Spots with very high molecular counts were filtered out for two of the samples, since these outliers did not seem to be dependent on tissue anatomy, but rather on technical artifacts (Extended Data Fig. 5a). Furthermore, spots with high mitochondrial content were not filtered out since they were located in a distinct region within each DG sample, that is to say the molecular layer (Extended Data Fig. 5b). This suggests that the mitochondrial counts are biologically relevant in each sample given the spatial pattern of the spots. The samples were then combined into a single Seurat object to allow us to perform analyses using the gene expression data from all samples (14 221 spots in total).

The following preprocessing steps were done using Seurat. The data was normalized using the log-normalization procedure with NormalizeData function. We next identified variable genes using the FindVariableFeatures function (with the selection method set to vst and the number of features set to 2000). The normalized dataset was next scaled using the ScaleData function and principal component analysis (PCA) was performed on the scaled counts by computing the top 30 principal components (PCs) with the RunPCA function. The first 15 PCs were used for downstream analysis. The merged Seurat object was converted into a SingleCellExperiment^52^ (v.1.16.0) object for batch correction with the R package Harmony^53^ (v.0.1.0) and for spot clustering.

The manual annotations were exported and integrated in the merged Seurat object for a comparison between the clustering performance of four different clustering algorithms: kmeans with the R package stats^50^, mclust with the R package mclust (v. 5.4.7)^54^, BayesSpace normal error method, and BayesSpace t-distributed error method from BayesSpace^15^. The clustering resolution was set at k=7 for each algorithm as there were seven manually annotated clusters in each section (see Manual annotation). The clustering accuracy for each algorithm was assessed using the adjusted rand index, as previously done in Zhao et al. (2021)^15^ and Maynard et al. (2021)^39^. Based on the ARI, unsupervised clustering was performed using the BayesSpace (t-distributed error model) clustering algorithm (v. 1.4.1) with the SpatialCluster (nrep = 50000) BayesSpace function, (Extended Data Fig. 1). Seurat was used for clustering visualization using the SpatialPlot function. We assessed the average expression of different canonical cell type markers in each BayesSpace cluster from all sections by generating a dotplot with the DotPlot function from Seurat. The following well-known cell type markers were used for dotplot visualization: *SATB2, SLC17A7, SLC17A6* (excitatory neurons); *GAD1, GAD2* (inhibitory neurons); *TMEM119, CX3CR1, MRC1* (microglia); *CLDN5, VTN* (endothelial cells); *GLUL, SOX9, AQP4, GFAP, ALDH1L1, VIM* (astrocytes); *PDGFRA, OLIG1, OLIG2* (oligodendrocyte precursor cells); *PLP1, MOG, MOBP, MBP* (oligodendrocytes); *SNAP25, STMN2, RBFOX3* (neurons).

### Visium sub-spot level analysis

We next used the SpatialEnhance (nrep = 200 000) and EnhanceFeatures functions of BayesSpace to computationally map the neurogenic features at the sub-spot level, where each Visium spot was divided into six sub-spots with BayesSpace. While Visium spots may contain up to 30 cells due to their 55-μm diameter, each of the generated sub-spots represented the transcriptomic signatures of a lower number of cells, thus increasing the cellular resolution of the data. The default nonlinear regression model (XGBoost) was trained for each neurogenic marker to determine the gene expression measured at the spot-level. The fitted model was next used to measure gene expression with the enhanced clustering. The spatial feature plots were generated using the featurePlot function from BayesSpace.

For the sub-spot level analysis of the Maynard et al. (2021)^39^, the publicly available dataset of the dorsolateral prefrontal cortex spatial transcriptomic data was downloaded, preprocessed, batch-corrected and computed at enhanced resolution with the integrated workflow of BayesSpace (v. 1.4.1). Four samples from the study were included in the analysis: samples 151671, 151672, 151673, 151674 (2M/2F, mean age=38.41 years old).

### Spot deconvolution with single-nucleus RNA-sequencing

We downloaded the publicly available human hippocampal dataset from Habib et al., (2017)^12^ (https://www.gtexportal.org/home/datasets). Spot deconvolution was performed using the R package SPOTlight’s seeded non-negative matrix factorization regression model^19^ (v. 0.1.7), which determined the topic profiles, in other words the set of defining genes, within each cell-type from the single-nucleus RNA sequencing dataset and within each capture spot. The marker genes in each cluster from the Habib et al. (2017) dataset were found using the FindAllMarkers function (with Wilcoxon Rank Sum test) with Seurat. The spotlight_deconvolution function with the standard parameters from SPOTlight was applied to determine the topic profiles from the single-nucleus dataset that fit best each spot’s topic profile. The deconvolution was next assessed by looking at how specific the topic profiles were for each cell type from the single-nucleus RNA sequencing dataset (Extended Data Fig. 6). The SpatialFeaturePlot function from Seurat was used to view the predicted cell-type proportions of the exDG (granule cell) and NSC (neural stem cell) clusters on the spatial data.

### RNAscope and data quantification

Frozen dentate gyrus samples from the 2 year old and 16 year old subjects, as well as the those used for the spatial transcriptomic experiments were cryosectioned at 10 μm, collected on Superfrost charged slides, and stored at −80°C. *In situ* hybridization was performed using Advanced Cell Diagnostics RNAscope® probes and reagents (ACD Bio, RNAscope® Multiplex Fluorescent v2 Assay) following the manufacturer’s instructions. Briefly, tissue sections were fixed in 10% neutral buffered formalin for 15 minutes at 4°C. A series of dehydration steps was next performed with different concentrations of ethanol baths (50%, 70%, 95%, 100%) and the sections were air dried for 5 minutes. A hydrogen peroxide treatment (10 minutes at room temperature) and a protease digestion (Protease Plus 1/15 dilution; 30 minutes at room temperature) were then done on the tissue sections. Sections were incubated for two hours in a temperature-controlled oven (HybEZ II, ACDbio) with different probe combinations: (1) Hs-GAD1 (catalog no. 404031), Hs-DCX-C3 (catalog no. 489551); (2) Hs-SLC17A7 (catalog no. 415611), Hs-DCX-C3; (3) Hs-NEUROD1 (catalog no. 437281), Hs-DCX-C3; (4) Hs-PROX1 (catalog no. 530241), Hs-CALB2-C2 (catalog no. 422171-C2), Hs-DCX-C3; (5) Hs-STMN1-O1 (catalog no. 471021), Hs-PROX1-C2 (catalog no. 530241-C2), Hs-DCX-C3; (6) Hs-NES (catalog no. 486341), Hs-ALDH1L1-C2 (catalog no. 438881-C2), Hs-SOX2-C3 (catalog no. 400871-C3); (7) Hs-PCNA (catalog no. 553071), Hs-PROX1-C2, Hs-DCX-C3; (8) Hs-MCM2 (catalog no. 451461), Hs-PROX1-C2, Hs-DCX-C3; (9) Hs-TMEM119 (catalog no. 478911), Hs-PROX1-C2, Hs-DCX-C3; (10) Hs-PROX1, Hs-ALDH1L1-C2, Hs-DCX-C3; and (11) Hs-MBP (catalog no. 411051), Hs-PROX1-C2, Hs-DCX-C3. After amplification steps (AMP1–3), probes were fluorescently labeled with Opal Dyes (Opal 520 product no. FP1487001KT, Opal 570 product no. FP1488001KT, Opal 690 product no. FP1497001KT; Perkin Elmer). Autofluorescence from lipofuscin was quenched using the reagent TrueBlack (Biotium) for 40 seconds and sections were finally coverslipped with Vectashield mounting medium containing 4′,6-diamidino-2-phenylindole (DAPI) to label the nucleus (Vector Laboratories, H-1800). Slides were kept at 4 °C until imaging. Both positive and negative controls supplied by the manufacturer were used on separate sections to verify signal specificity. For frozen-fixed samples, two additional steps were performed: a 30-minute incubation step before formalin fixation and an antigen retrieval step before a 30-minute protease digestion at 40°C (Protease IV).

The slides were imaged at 20X magnification using the Olympus VS120 virtual slide scanner and the scans were transferred to QuPath^55^ (v.0.3.0) for further analysis. Using QuPath’s automated cell detection based on DAPI staining and manual counting, the percentage of cells positive for different combinations of neurogenic markers was counted in two layers of interest: the GCL and SGZ. For each section, the GCL was traced based on the DAPI staining, demonstrating a tightly compact layer of DG granule neurons, and the SGZ was defined as a 50 μm region below the GCL, as previously done in Seki et al. (2019)^6^. For stringent quantification, cells with 4 or more fluorescent puncta were counted as positive for *GAD1*, *NEUROD1*, *CALB2*, *NES*, *SOX2*, *TMEM119*, *ALDH1L1*, *MBP*, *PCNA*, *MCM2*, and *DCX* probes and cells with at least 6 fluorescent puncta were labeled as positive for *SLC17A7*, *PROX1*, *STMN1*. The data was plotted using the ggplot2 (v. 3.3.5) R package^56^ and in GraphPad (version 7, GraphPad Software LLC).

### Image acquisition

Images included in the figures were taken on a FV1200 laser scanning confocal microscope using a 40x objective (NA: 0.95). The XY axis pixel number (1600×1600), Kalman averaging (2), and laser scanning speed (2µs/pixel) were modified to these settings to improve image resolution. Laser power and detection voltage parameters were adjusted between subjects for each set of experiments to improve image quality. All parameters were optimized to decrease signal to noise ratio and remove autofluorescence from lipofuscin and cellular debris.

### Statistical analysis

Statistical analyses were performed using GraphPad (version 7) software. The two-sided Pearsons’s correlation test was used to examine the association between dependent variables and covariates, such as age and PMI. The graphs generated from these analyses represent either mean values±SD or values from one staining replicate per probe combination from whole DG sections. A 95% confidence interval was used for statistical comparisons.

## Acknowledgments

The authors are grateful to Dominique Mirault, Jean-François Théroux and Zhipeng Niu for technical assistance, and to Dr. Nada Jabado for providing one of the brain samples used in this study. This work was funded by a Healthy Brains, Healthy Lives (HBHL; CFREF) Innovative Ideas grant to N.M. R.R. received postdoctoral fellowships from FRQ-S and AMH. The Douglas-Bell Canada Brain Bank is funded by platform support grants to G.T. and N.M. from the RQSHA (FRQ-S), HBHL, and Brain Canada.

## Author information

### Authors and Affiliations

**McGill Group for Suicide Studies, Douglas Mental Health University Institute, McGill University, Montreal, Canada**

Sophie Simard, Reza Rahimian, Maria Antonietta Davoli, Stéphanie Théberge, Gustavo Turecki, Corina Nagy & Naguib Mechawar

**Illawarra Health and Medical Research Institute, University of Wollongong, Wollongong, Australia.**

Natalie Matosin

**School of Chemistry and Molecular Bioscience, Faculty of Science, Medicine and Health, University of Wollongong, Wollongong, Australia.**

Natalie Matosin

**Department of Psychiatry, McGill University, Montréal, Canada**

Gustavo Turecki, Corina Nagy & Naguib Mechawar

### Contributions

G.T., C.N. and N.M. conceptualized the study. C.N. and M.A.D designed and performed the Visium Spatial Gene Expression experiments. S.S. analyzed the spatial transcriptomic data. S.S., R.R., and M.A.D. designed the fluorescent *in situ* hybridization experiments. S.S. performed and analyzed the fluorescent *in situ* hybridization experiments. S.T. performed confocal imaging. S.S. and N.M wrote the first draft of the manuscript which was revised and edited by all authors. C.N. and N.M. supervised the study.

### Corresponding author

Correspondence to Naguib Mechawar, Ph.D.

## FIGURE LEGENDS

**Extended Data Figure 1. Clustering accuracy comparison for unsupervised clustering (related to Figure 1).**

**(a)** Manual annotations created on Loupe Browser and plotted on each section with Seurat. **(b)** Clustering accuracy comparison between four clustering algorithms (kmeans, mclust, BayesSpace normal error model and BayesSpace t-distributed error model) with manual annotations used as a point of reference. The adjusted rand index (ARI) was used as a measure of comparison. **(c)** Clustering of each tested algorithm spatially plotted on each section with the clustering resolution set at k=7.

**Extended Data Figure 2. Correlation between the number of *DCX*^+^ cells per mm^2^, age and PMI (related to Figure 2 and Methods: Human post-mortem tissue)**

**(a)** Correlation between the average number of *DCX*^+^ cells per mm^2^ and age in the SGZ (two-sided Pearson’s correlation, r=0.03127, P=0.9414) and GCL (two-sided Pearson’s correlation, r=- 0.7517, P=0.0315). The graph shows mean values ± SD from all subjects with four staining replicates per subject. A significant negative correlation between these parameters was observed in the GCL. **(b)** Correlation between the average number of *DCX*^+^ cells per mm^2^ and post-mortem interval (PMI) in the SGZ (two-sided Pearson’s correlation, r=0.5669, P=0.1845) and GCL (two-sided Pearson’s correlation, r=0.6872, P=0.0881). The graph shows mean values ± SD from all subjects, except the infant (due to unavailability of information), with four staining replicates per subject. No significant correlation between these parameters was observed. **(c)** Correlation between the percentage of *DCX*^+^ cells expressing *SLC17A7* (two-sided Pearson’s correlation, r=- 0.8661, P=0.0054), *GAD1* (two-sided Pearson’s correlation, r=0.02026, P=0.9620), *NEUROD1* (two-sided Pearson’s correlation, r=-0.9454, P=0.0004), or *PROX1* (two-sided Pearson’s correlation, r=-0.6875, P=0.0595), and age in the GCL. For each subject, there is one staining replicate per probe combination. A statistically significant correlation was observed between age and the percentage of *DCX*^+^*SLC17A7*^+^ and *DCX*^+^*NEUROD1*^+^ cells. **(d)** Correlation between the percentage of *DCX*^+^ expressing *SLC17A7* (two-sided Pearson’s correlation, r=-0.8991, P=0.0024), *GAD1* (two-sided Pearson’s correlation, r=0.5872, P=0.1259), *NEUROD1* (two-sided Pearson’s correlation, r=-0.7327, P=0.0387), and *PROX1* (two-sided Pearson’s correlation, r=-0.9624, P=0.0001), and age in the SGZ. For each subject, there is one staining replicate per probe combination. SGZ, subgranular zone; GCL, granule cell layer. *0.05 >P≥ 0.01; **0.01 >P≥ 0.001; ***P< 0.001 in two-sided Pearson’s correlation. The asterisks indicate statistical significance in Pearson’s correlation.

**Extended Data Figure 3. *DCX* expression in cells expressing glial markers.**

**(a)** *DCX* expression in cells positive for *TMEM119* (microglia) and **(b)** *ALDH1L1* (astrocytes) in the DG of 26 year old and 42 year old subjects respectively. Scale bars = 20 μm (scale bars for expanded insets = 10 μm).

**Extended Data Figure 4. *STMN1* expression in the human infant and adult DG (related to Figure 2).**

**(a)** *STMN1* expression in *PROX1*^+^*DCX*^+^ cells located in the SGZ and GCL of the adult human DG. Scale bars = 20 μm. **(b)** Percentage of *PROX1*^+^*DCX^+^* expressing *STMN1* in the SGZ and GCL of the whole DG in the infant and adult dentate gyrus. **(c)** Log-normalized expression of *STMN1* spatially plotted onto each section (n=4) at the sub-spot level using BayesSpace. Higher values in the scale correspond to higher expression levels, whereas lower values reflect lower expression levels. SGZ, subgranular zone; GCL, granule cell layer.

**Extended Data Figure 5. Quality check steps for unsupervised clustering (related to Methods: Clustering accuracy comparison and unsupervised spot clustering).**

**(a)** Spatial representation of the molecular counts in two of the sections, showing outlier spots with higher than 60 000 UMIs (unique molecular identifiers) and 30 000 UMIs respectively. These outlier spots were filtered out from the spatial data. The spatial distribution of the molecular counts is plotted with Seurat. **(b)** Spots with high percentages of mitochondrial gene expression (percent.mt) are located within the region of the molecular layer in each section. The coloured scale corresponds to the percentage of mitochondrial genes in each spot. The spatial distribution of mitochondrial reads is plotted on each section with Seurat.

**Extended Data Figure 6. Topic profiles for each cluster from Habib et al., (2017) with the R package SPOTlight (related to Methods: Spot deconvolution with single-nucleus RNA-sequencing).**

**(a)** Specificity of each topic profile detected in Habib *et al*., (2017) with SPOTlight’s seeded non-negative matrix factorization (NMF) regression model. The size of dots represents the proportion of nuclei from each cluster corresponding to each topic profile. **(b)** Individual topic profiles of each nucleus from the exDG and NSC clusters, with nuclei showing a similar distribution within each cluster.

**Supplementary Table 1. Subject information and sequencing metrics obtained with Space Ranger of postmortem human hippocampal samples used for Visium Spatial Gene Expression (related to** Figure 1 **and Methods: Human postmortem tissue).**

**Supplementary Table 2. Percentage of spots expressing canonical cell type markers (based on scaled average expression) in each spot-level BayesSpace cluster (related to Figure 1).**

**Supplementary Table 3. Subject information of postmortem human hippocampal samples used for RNAscope (related to Figures 2-3 and Methods: Human postmortem tissue).**

## References

1. Eriksson, P.S., et al. Neurogenesis in the adult human hippocampus. Nat. Med. 4, 1313–1317 (1998).

2. Boldrini, M., et al. Human hippocampal neurogenesis persists throughout aging. Cell Stem Cell 22, 589–599.e585 (2018).

3. Dennis, C.V., Suh, L.S., Rodriguez, M.L., Kril, J.J. & Sutherland, G.T. Human adult neurogenesis across the ages: An immunohistochemical study. Neuropathol. Appl. Neurobiol. 42, 621–638 (2016).

4. Flor-García, M., et al. Unraveling human adult hippocampal neurogenesis. Nat. Protoc. 15, 668–693 (2020).

5. Moreno-Jiménez, E.P., et al. Adult hippocampal neurogenesis is abundant in neurologically healthy subjects and drops sharply in patients with Alzheimer’s disease. Nat. Med. 25, 554–560 (2019).

6. Seki, T., Hori, T., Miyata, H., Maehara, M. & Namba, T. Analysis of proliferating neuronal progenitors and immature neurons in the human hippocampus surgically removed from control and epileptic patients. Sci. Rep. 9, 18194 (2019).

7. Sorrells, S.F., et al. Human hippocampal neurogenesis drops sharply in children to undetectable levels in adults. Nature 555, 377–381 (2018).

8. Terreros-Roncal, J., et al. Impact of neurodegenerative diseases on human adult hippocampal neurogenesis. Science 374, 1106–1113 (2021).

9. Tobin, M.K., et al. Human hippocampal neurogenesis persists in aged adults and Alzheimer’s disease patients. Cell Stem Cell 24, 974–982.e973 (2019).

10. Ayhan, F., et al. Resolving cellular and molecular diversity along the hippocampal anterior-to-posterior axis in humans. Neuron 109, 2091–2105.e2096 (2021).

11. Franjic, D., et al. Transcriptomic taxonomy and neurogenic trajectories of adult human, macaque, and pig hippocampal and entorhinal cells. Neuron 110, 452–469.e414 (2022).

12. Habib, N., et al. Massively parallel single-nucleus RNA-seq with DroNc-seq. Nat Methods 14, 955–958 (2017).

13. Wang, W., et al. Transcriptome dynamics of hippocampal neurogenesis in macaques across the lifespan and aged humans. Cell Res. 32, 729–743 (2022).

14. Zhou, Y., et al. Molecular landscapes of human hippocampal immature neurons across lifespan. Nature 607, 527–533 (2022).

15. Zhao, E., et al. Spatial transcriptomics at subspot resolution with BayesSpace. Nat. Biotechnol. 39, 1375–1384 (2021).

16. Iwano, T., Masuda, A., Kiyonari, H., Enomoto, H. & Matsuzaki, F. Prox1 postmitotically defines dentate gyrus cells by specifying granule cell identity over CA3 pyramidal cell fate in the hippocampus. Development 139, 3051–3062 (2012).

17. Gao, Z., et al. Neurod1 is essential for the survival and maturation of adult-born neurons. Nat. Neurosci. 12, 1090–1092 (2009).

18. von Bohlen und Halbach, O. Immunohistological markers for proliferative events, gliogenesis, and neurogenesis within the adult hippocampus. Cell Tissue Res. 345, 1–19 (2011).

19. Elosua-Bayes, M., Nieto, P., Mereu, E., Gut, I. & Heyn, H. SPOTlight: seeded NMF regression to deconvolute spatial transcriptomics spots with single-cell transcriptomes. Nucleic Acids Res. 49, e50 (2021).

20. Sorrells, S.F., et al. Positive controls in adults and children support that very few, if any, new neurons are born in the adult human hippocampus. J. Neurosci. 41, 2554–2565 (2021).

21. Kelley, K.W., Nakao-Inoue, H., Molofsky, A.V. & Oldham, M.C. Variation among intact tissue samples reveals the core transcriptional features of human CNS cell classes. Nat. Neurosci. 21, 1171–1184 (2018).

22. Terstege, D.J., Addo-Osafo, K., Campbell Teskey, G. & Epp, J.R. New neurons in old brains: implications of age in the analysis of neurogenesis in post-mortem tissue. Mol. Brain 15, 38 (2022).

23. Seki, T. Understanding the real state of human adult hippocampal neurogenesis from studies of rodents and non-human primates. Front. Neurosci. 14, 839 (2020).

24. Lucassen, P.J., et al. Limits to human neurogenesis-really? Mol. Psychiatry 25, 2207–2209 (2020).

25. Snyder, J.S. Recalibrating the relevance of adult neurogenesis. Trends Neurosci. 42, 164–178 (2019).

26. Roybon, L., et al. Neurogenin2 directs granule neuroblast production and amplification while NeuroD1 specifies neuronal fate during hippocampal neurogenesis. PLoS One 4, e4779 (2009).

27. Walker TL., et al. The doublecortin-expressing population in the developing and adult brain contains multipotential precursors in addition to neuronal-lineage cells. J Neurosci. 273734–3742 (2007).

28. Boulanger JJ & Messier C. Doublecortin in Oligodendrocyte Precursor Cells in the Adult Mouse Brain. Front Neurosci. 11 (2017).

29. Verwer, R.W., et al. Mature astrocytes in the adult human neocortex express the early neuronal marker doublecortin. Brain 130, 3321–3335 (2007).

30. Brandt, M.D., et al. Transient calretinin expression defines early postmitotic step of neuronal differentiation in adult hippocampal neurogenesis of mice. Mol. Cell. Neurosci. 24, 603–613 (2003).

31. Lavado, A., Lagutin, O.V., Chow, L.M., Baker, S.J. & Oliver, G. Prox1 is required for granule cell maturation and intermediate progenitor maintenance during brain neurogenesis. PLoS Biol. 8 (2010).

32. Kohler, S.J., Williams, N.I., Stanton, G.B., Cameron, J.L. & Greenough, W.T. Maturation time of new granule cells in the dentate gyrus of adult macaque monkeys exceeds six months. Proc. Natl. Acad. Sci. U. S. A. 108, 10326–10331 (2011).

33. La Rosa, C., Ghibaudi, M. & Bonfanti, L. Newly generated and non-newly generated "immature" neurons in the mammalian brain: a possible reservoir of young cells to prevent brain aging and disease? J Clin Med 8 (2019).

34. Ngwenya, L.B., Peters, A. & Rosene, D.L. Maturational sequence of newly generated neurons in the dentate gyrus of the young adult rhesus monkey. J. Comp. Neurol. 498, 204–216 (2006).

35. Hagihara, H., et al. Expression of progenitor cell/immature neuron markers does not present definitive evidence for adult neurogenesis. Mol. Brain 12, 108 (2019).

36. Li, Y.N., et al. Doublecortin-expressing neurons in human cerebral cortex layer ii and amygdala from infancy to 100 years old. Mol. Neurobiol. (2023).

37. Maheu ME, Davoli MA, Turecki G, Mechawar N. Amygdalar expression of proteins associated with neuroplasticity in major depression and suicide. J Psychiatr Res. 47, 384–390 (2013). doi:10.1016/j.jpsychires.2012.11.013

38. Sorrells, S.F., et al. Immature excitatory neurons develop during adolescence in the human amygdala. Nat Commun 10, 2748 (2019).

39. Maynard, K.R., et al. Transcriptome-scale spatial gene expression in the human dorsolateral prefrontal cortex. Nat. Neurosci. 24, 425–436 (2021).

40. Duque, A., Arellano, J.I. & Rakic, P. An assessment of the existence of adult neurogenesis in humans and value of its rodent models for neuropsychiatric diseases. Mol. Psychiatry 27, 377–382 (2022).

41. Augusto-Oliveira, M., Arrifano, G.P.F., Malva, J.O. & Crespo-Lopez, M.E. Adult hippocampal neurogenesis in different taxonomic groups: Possible functional similarities and striking controversies. Cells 8 (2019).

42. Kempermann, G. The neurogenic reserve hypothesis: what is adult hippocampal neurogenesis good for? Trends Neurosci. 31, 163–169 (2008).

43. Kempermann, G., et al. Human adult neurogenesis: evidence and remaining questions. Cell Stem Cell 23, 25–30 (2018).

44. Merkley, C.M., Jian, C., Mosa, A., Tan, Y.F. & Wojtowicz, J.M. Homeostatic regulation of adult hippocampal neurogenesis in aging rats: long-term effects of early exercise. Front. Neurosci. 8, 174 (2014).

45. Shevtsova, O., Tan, Y.F., Merkley, C.M., Winocur, G. & Wojtowicz, J.M. Early-Age Running Enhances Activity of Adult-Born Dentate Granule Neurons Following Learning in Rats. eNeuro 4 (2017).

46. Schwab, C., Yu, S., Wong, W., McGeer, E.G. & McGeer, P.L. GAD65, GAD67, and GABAT immunostaining in human brain and apparent GAD65 loss in Alzheimer’s disease. J. Alzheimers Dis. 33, 1073-1088 (2013).

47. Cabezas, C., Irinopoulou, T., Cauli, B. & Poncer, J.C. Molecular and functional characterization of GAD67-expressing, newborn granule cells in mouse dentate gyrus. Front Neural Circuits 7, 60 (2013).

48. Zhao, S., et al. Fluorescent labeling of newborn dentate granule cells in GAD67-GFP transgenic mice: a genetic tool for the study of adult neurogenesis. PLoS One 5 (2010).

49. Knoth, R., et al. Murine features of neurogenesis in the human hippocampus across the lifespan from 0 to 100 years. PLoS One 5, e8809 (2010).

50. Team, R.C. R: A language and environment for statistical computing. R Foundation for Statistical Computing, Vienna, Austria. (2021).

51. Stuart, T., et al. Comprehensive integration of single-cell data. Cell 177, 1888–1902.e1821 (2019).

52. Amezquita R., et al. Orchestrating single-cell analysis with Bioconductor. Nature Methods 17, 137–145 (2020).

53. Korsunsky, I., et al. Fast, sensitive and accurate integration of single-cell data with Harmony. Nat Methods 16, 1289–1296 (2019).

54. Scrucca, L., Fop, M., Murphy, T.B. & Raftery, A.E. mclust 5: clustering, classification and density estimation using gaussian finite mixture models. R j 8, 289–317 (2016).

55. Bankhead, P., et al. QuPath: Open source software for digital pathology image analysis. Sci. Rep. 7, 16878 (2017).

56. Wickham, H. ggplot2: elegant graphics for data analysis, Springer-Verlag New York (2016).

