## Supplementary figures and images for "Spatial transcriptomic analysis of adult hippocampal neurogenesis in the human brain"

### Extended Data Figure 1

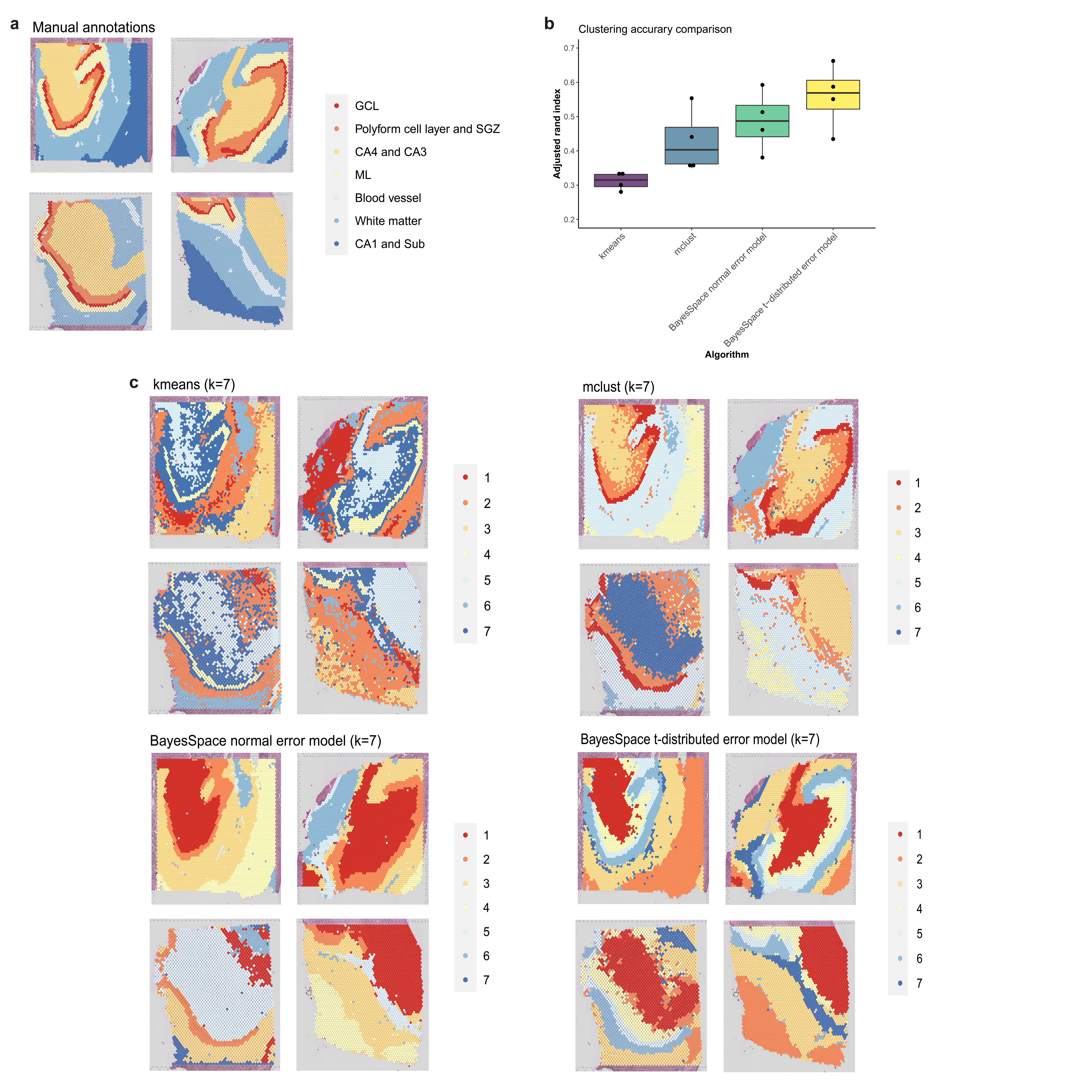

### Extended Data Figure 2

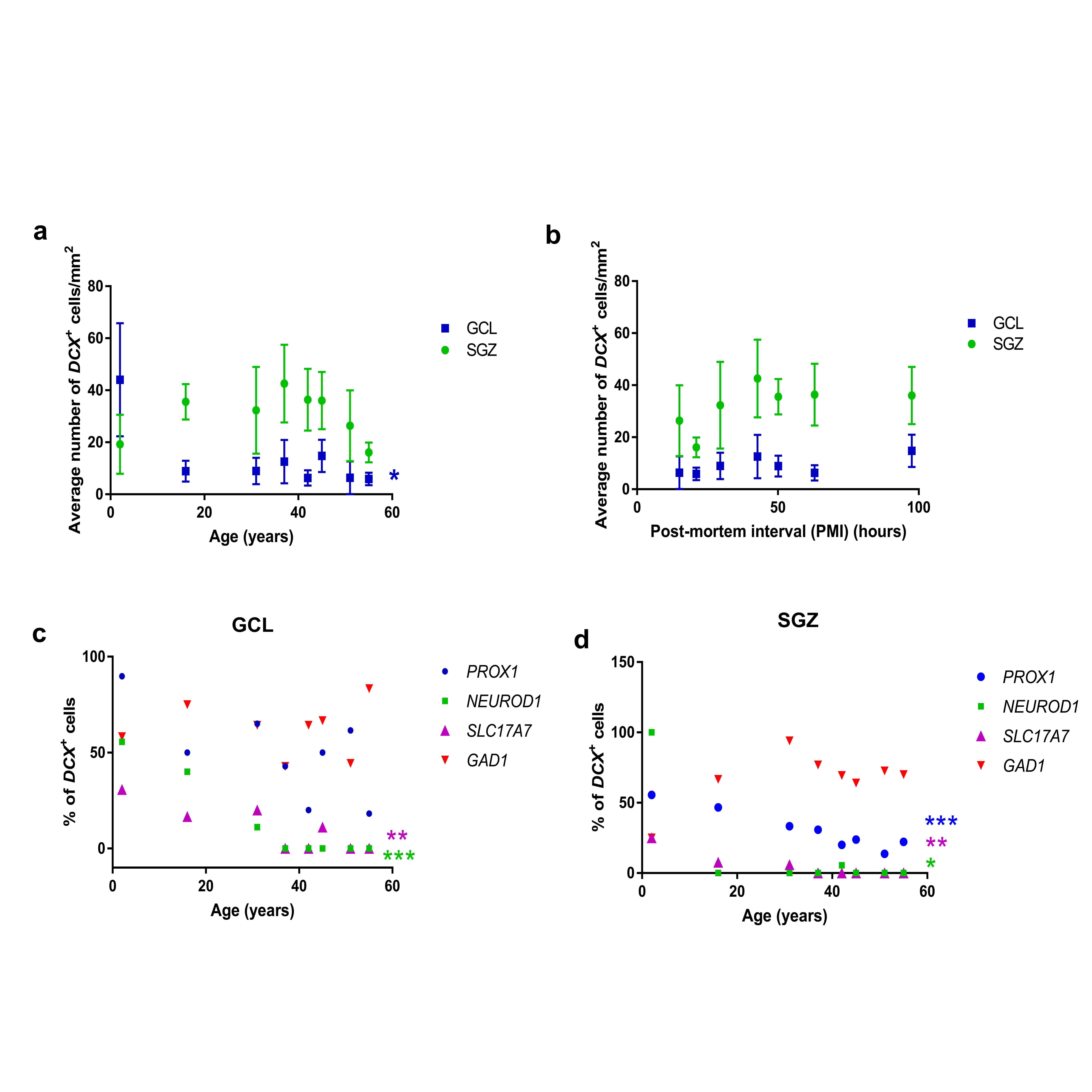

### Extended Data Figure 3

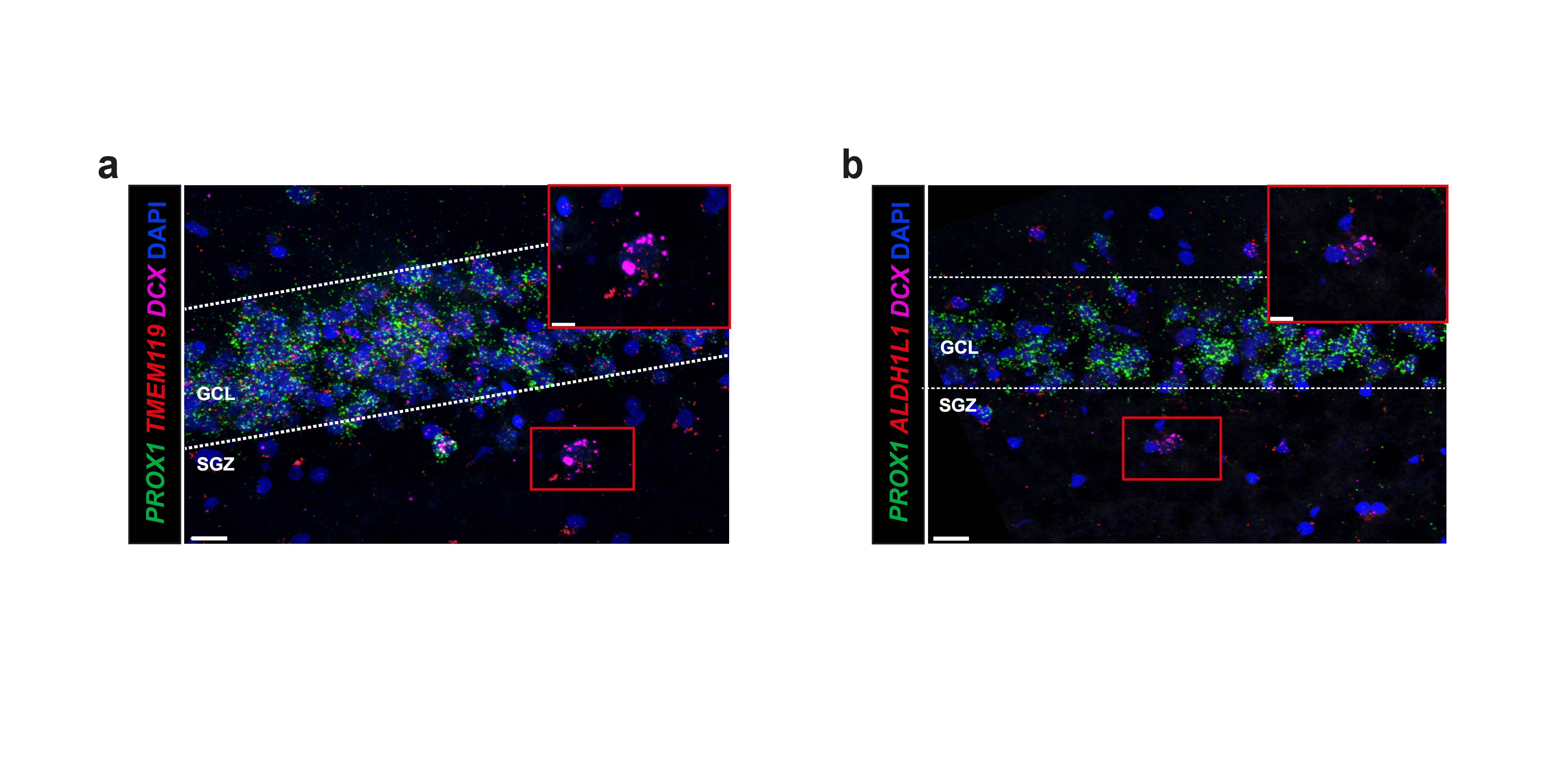

### Extended Data Figure 4

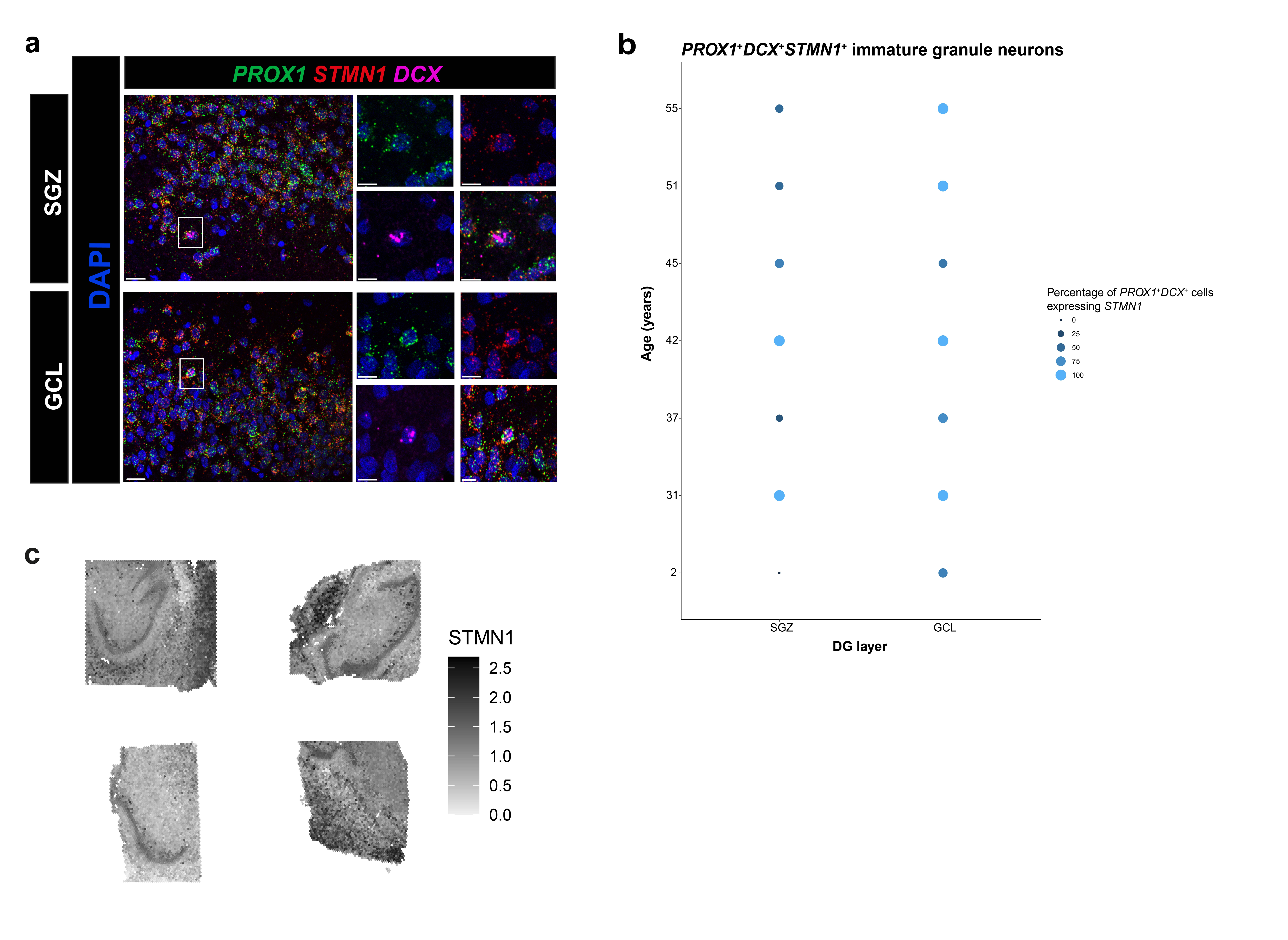

### Extended Data Figure 5

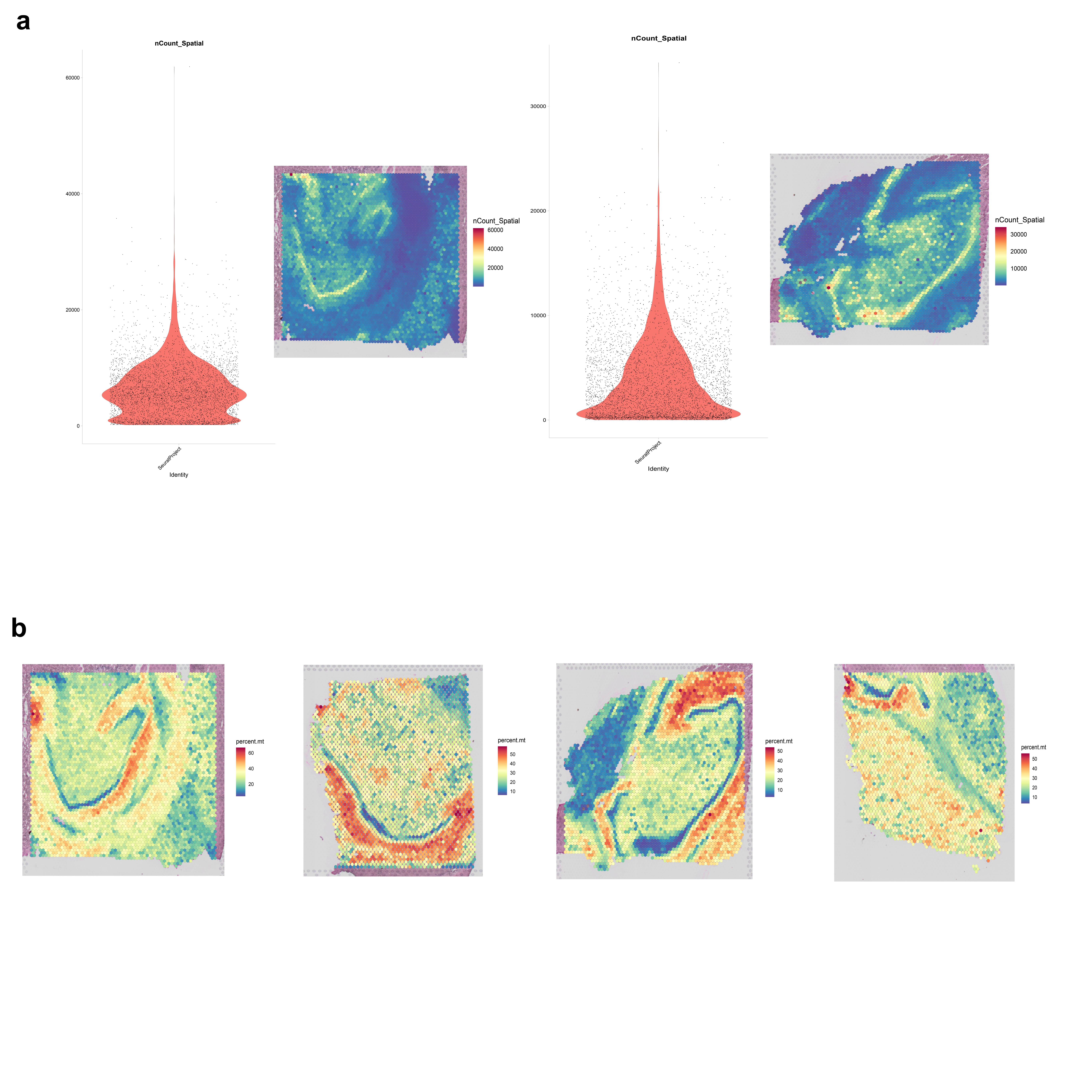

### Extended Data Figure 6

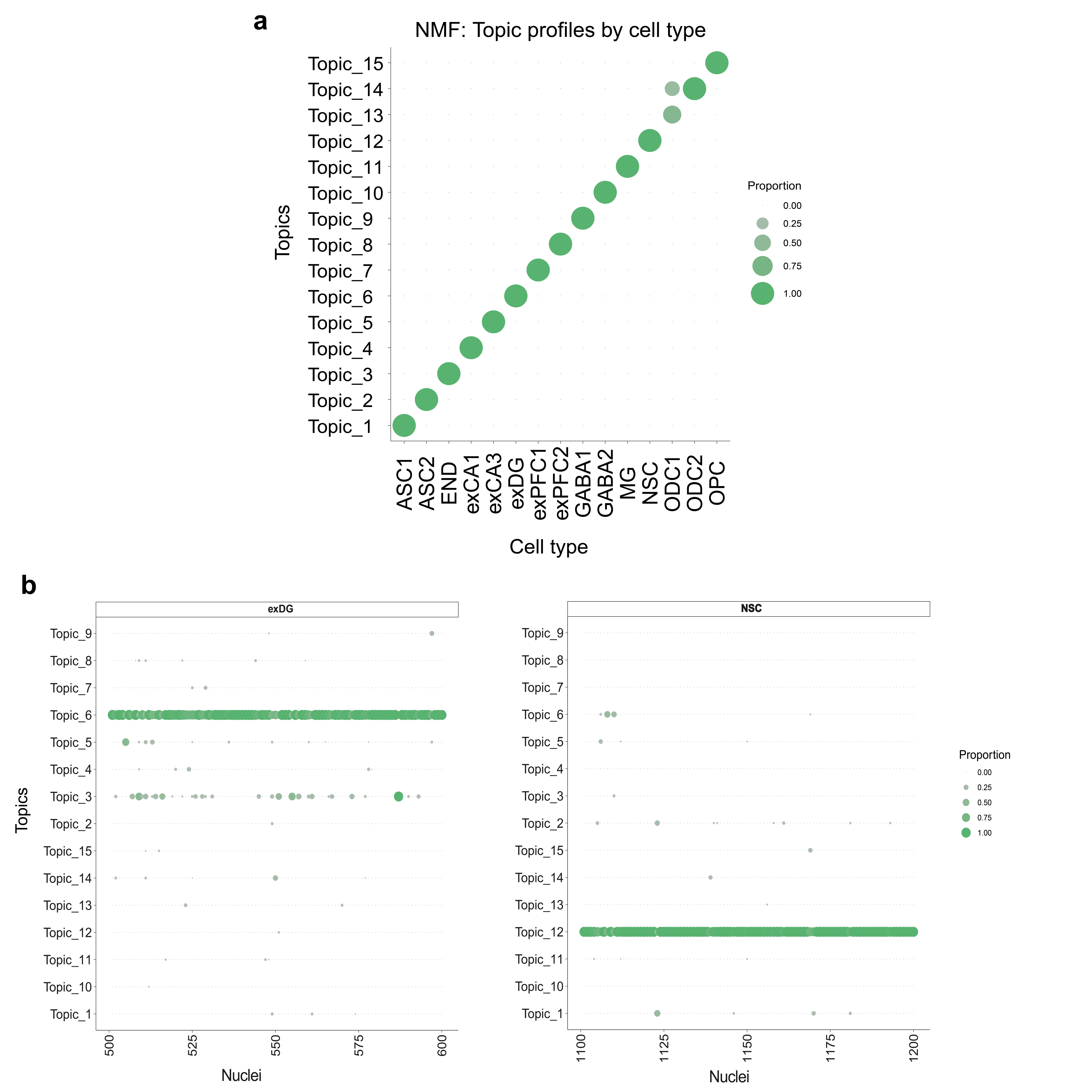
